# Locus-specific introgression in young hybrid swarms: drift dominates selection

**DOI:** 10.1101/2020.09.17.300434

**Authors:** S. Eryn McFarlane, Helen V. Senn, Stephanie L. Smith, Josephine M. Pemberton

## Abstract

Closely related species that have previously inhabited geographically separated ranges are hybridizing at an increasing rate due to human disruptions. These anthropogenic hybrid zones can be used to study reproductive isolation between species at secondary contact, including examining locus-specific rates of introgression. Introgression is expected to be heterogenous across the genome, reflecting variation in selection. Those loci that introgress especially slowly are good candidates for being involved in reproductive isolation, while those loci that introgress quickly may be involved in adaptive introgression. In the context of conservation, policy makers are especially concerned about introduced alleles moving quickly into the background of a native or endemic species, as these alleles could replace the native alleles in the population, leading to extinction via hybridization. We applied genomic cline analyses to 44997 SNPs to identify loci introgressing at excessive rates when compared to the genome wide expectation in an anthropogenic hybridizing population of red deer and sika in Kintyre Scotland. We found 11.4% of SNPs had cline centers that were significantly different from the genome wide expectation, and 17.6% had excessive rates of introgression. Based on simulations, we believe that many of these markers have diverged from average due to drift, rather than because of selection. Future work could determine the policy implications of allelic-replacement due to drift rather than selection, and could use replicate, geographically distinct hybrid zones to narrow down those loci that are indeed responding to selection in anthropogenic hybrid zones.

## Introduction

The rate of hybridization between closely related species that have recently come into secondary contact is increasing, due to increased human-assisted migration and environmental change (Parmesan and Yohe 2003, Grabenstein and Taylor 2018). While hybridization is not necessarily negative (Hamilton and Miller 2016), in many cases hybridization can cause problems for native species. If F1s are inviable or sterile then hybridization is a loss of reproductive effort (Allendorf et al. 2001). In contrast, the presence of viable, fertile hybrid offspring can lead to populations with large numbers of hybrids, and in the most extreme cases, whole populations comprised only of hybrid individuals (Allendorf et al. 2001). Biodiversity can be lost through hybridization, either if all remaining members of a species are hybrids (extinction via hybridization; Allendorf et al. 2001, Todesco et al. 2016, Allendorf and Luikart 2009, Rhymer and Simberloff 1996), or if particular endemic alleles are replaced by novel alleles introduced by backcrossing and driven to fixation via selection (as described by Petit 2004).

Hybrid zones, whether naturally occurring or due to human interference, can be used as ‘natural laboratories’ for research into selection and the genetics of reproductive isolation between species (Hewitt 1988). The rate of introgression of alleles between species is expected to be heterogenous across the genome, reflecting variation in selection (Baack and Rieseberg 2007). Backcrossing coupled with recombination will separate haplotypes that are commonly found together and create novel haplotypes where selection can act on alleles in unique genetic backgrounds (Arnold et al. 1999). Alleles that move quickly across the species barrier are assumed to be under positive selection in their new genetic background, while alleles that do not introgress between species are candidates for contributing to reproductive isolation (Baack and Rieseberg 2007). Drift will also be acting on these alleles, particularly if hybridization is rare or one of the parental populations is small. In these cases, we expect substantial variation in the degree of introgression across loci, as a result of the sampling error introduced by reproduction and recombination (Baird, Barton, and Etheridge 2003). If non-native alleles are increasing in frequency, whether due to selection or drift, we should apply the precautionary principle until we can be sure that selection will not bring these alleles to fixation. Identifying those endemic loci that are most likely to be replaced by novel alleles gives a target for policy makers to reflect upon and consider protecting.

Geographic cline analyses have been used to determine the extent of hybridization between two species at a contact zone (Barton and Hewitt 1985, Barton and Gale 1993). Traditionally, the width of these geographic gradients of allele frequencies can be used to infer selection on each allele as it introgresses from one species to another across a landscape (Mallet et al. 1990). Recently, *genomic* clines, which replace geographic gradients with hybrid indices, have been used in the same way, and have the advantage that they can be applied even when hybrids have a mosaic distribution, or in a hybrid swarm (Gompert and Buerkle 2012, Lexer et al. 2007, Gompert and Buerkle 2011). Genomic clines use a multinomial regression that predicts the probability of a particular genotype (*θ*) given a hybrid index (h), where:

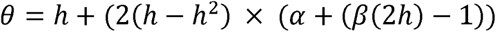

Here, *α* is analogous to the location of the cline center and can be interpreted as the direction of introgression, i.e. a positive *α* means excess ancestry from species A to species B and negative *α* means excess ancestry from species B to A. *β* is analogous to the width of the cline and can be interpreted as the strength of the barrier to gene flow (Janoušek et al. 2015). Positive *β* is interpreted as a narrow cline, where introgression is impeded, and negative *β* is a wide cline, where introgression is faster than expected based on the genomic expectation (Gompert and Buerkle 2009).

α and β are not explicitly expected to covary with each other (although they are not fully independent), nor are α and β necessarily expected to covary with divergence estimates between the parental species in the system such as Fst (Charlesworth 1998). However, those loci that are both highly diverged between species (i.e. high F_st_) and slow moving (large positive β) are good candidates for loci involved in reproductive isolation (Gompert and Buerkle 2009, Lexer et al. 2007), particularly if they are not expected to be highly diverged because of other genomic constraints (i.e. recombination cold spots; Burri et al. 2015, Cruickshank and Hahn 2014). Studies of naturally occurring hybridization regularly find many markers, spread across the genome, with significant α and β estimates, and typically find more loci that are significant for α than β loci (but see (Pulido-Santacruz, Aleixo, and Weir 2018) who found no divergent α or β SNPs between either *Willisornis* or *Xiphorhynchus* species pairs). For example, Janoušek et al. (2015) found that as many as 70% of SNPs diverged from genome-wide expections in a *mus* hybrid zone, Parchman et al. (2013) using 59 100 SNPs found more than 1000 significant α SNPs and more than 400 significant β SNPs between *Manacus candei* and *M. vinellinus*, and Sung et al. (2018) reported ∼30% of 45384 SNPs with significantly diverged α and ∼1% of SNPs with significantly diverged rates of β between *Iris hexagona* and *I. fulva*. The vast number of reported genome wide excess α and β SNPs from many systems are unlikely to all be related to selection, especially given that selection must be extremely strong to be detected at the genome-wide level in artificial selection studies (e.g. Castro et al. 2019). Simulations of admixed populations that varied population sizes found that, particularly with a population size of only 100, both α and β estimates could be quite variable, and when loci under selection were simulated, particularly when there was weak selection and low levels of admixture, there were high false discovery rates (Gompert and Buerkle 2011). Before genomic regions can be considered candidates to be responding to selection, careful consideration of expections due to non-selective forces must be undertaken (Gompert and Buerkle 2011).

The red deer (*Cervus elaphus*) is an emblematic animal native to Scotland. It was named as one of ‘Scotland’s big 5’ in a campaign to increase engagement with wildlife ran by Scottish government between 2013 and 2015 (Scottish Wildlife Trust, 2013), known for its large size, large antlers and bright red summer coat. Red deer are abundant through much of Scotland and they are popular for hunting (deer stalking) and with tourists and unpopular for their ecological impacts, particularly on young trees. Physically smaller Japanese sika (*C. nippon*) were introduced to Scotland in the late 19^th^ century, and have since hybridized with the red deer (Ratcliffe 1987). On the Kintyre peninsula, Argyll, more than 40% of sampled phenotypic red deer and sika individuals are hybrids according to 50 000 SNP markers, with the majority being the result of multiple generations of backcrossing (McFarlane et al. 2020). Hybrid deer tend towards an intermediate phenotype and thus are smaller, have smaller antlers, and are more likely to have the spots typical of sika than parental species red deer (Senn, Swanson, et al. 2010). While there is a trend from red deer in the north to sika in the south of the peninsula, the distribution of hybrids does not follow a cline, being instead concentrated in specific areas (Senn, Barton, et al. 2010). Additionally, in a study using 20 microsatellite markers, there was no evidence that the number of hybrid individuals was changing over a period of 15 years (Senn, Barton, et al. 2010).

In this study, we sought evidence among red-sika hybrids that specific genome regions have introgressed more or less than expected under neutrality, in ways that might be interpreted as being due to selection. We used 50K SNP genotypes in 222 Kintyre hybrid deer to estimate genomic clines and show that, as in the other studies cited above, many loci exceed background expectation in terms of direction of introgression α and cline width β. We then conduct population genetic simulations to investigate admixture scenarios that shed light on the likely roles of drift and selection in generating these results.

## Methods

### Sample Collection

513 deer samples were collected from 15 forestry sites in the Kintyre region of Scotland between 2006 and 2011. These samples were collected by the Forestry Commission Scotland (now Forestry and Land Scotland) as part of normal deer control measures. Deer were shot as encountered, without regard to the phenotype of the animal (Smith et al. 2018). Sample collection consisted of ear tissue and has been previously described elsewhere (Senn and Pemberton 2009, Smith et al. 2018). Samples were either preserved in 95% ethanol or frozen for long-term storage.

### DNA extraction and SNP Genotyping

We used the DNeasy Blood and Tissue Kit (Qiagen) according to the manufacturer’s instructions to extract DNA for SNP analysis, with the exception that we eluted twice in 25μl buffer TE to obtain DNA at a sufficiently high concentration. Concentration was assayed using the Qubit™ dsDNA BR Assay Kit (Invitrogen). Any samples below 50 ng/μl were vacuum-concentrated, re-extracted or omitted from SNP analysis.

SNPs were genotyped on the Cervine Illumina iSelect HD Custom BeadChip using an iScan instrument following manufacturer’s instructions (as in (Huisman et al. 2016). When this SNPchip was developed, SNPs were spaced evenly throughout the genome based on the bovine genome, with which the deer genome has high homology. We used a positive control twice on each 96 well plate to check for consistency between batches (Huisman et al. 2016). We scored genotypes using GenomeStudio using the clusters from Huisman et al (2016), and clustered SNPs manually if they could not be resolved in these clusters (McFarlane et al. 2020). All quality control was done in PLINK (Purcell et al. 2007). We excluded individual samples with a call rate of less than 0.90, and deleted loci with a minor allele frequency of less than 0.001 and/or a call rate of less than 0.90. We did not exclude SNPs based on Hardy Weinberg Equilibrium (HWE) as highly differentiated markers between red and sika are not expected to be in HWE. When the chip was designed, the majority of the 53K SNPs included were selected to be polymorphic in red deer, 4500 SNPs were selected to be diagnostic between either red deer and sika or red deer and wapiti (*Cervus canadensis*) (Brauning et al. 2015). Of these 629 SNPs are diagnostic and an additional 3205 SNPs are ancestry informative markers (hereafter together as AIMs) in Kintyre. These AIMs were determined based on having extreme allele frequency differences where the differences in frequency between the two populations was more than 0.95 (McFarlane et al. 2020). While one pool of 12 sika from Kintyre were whole genome sequenced for the development of this SNP chip, the focus was on polymorphic SNPs in red deer on Rum (Brauning et al. 2015). A high-density deer linkage map confirms high homology between cervine and bovine genomes (Johnston et al. 2017); in the present study we have used the bovine map as this allows use of all of the SNPs, including those that are not polymorphic in red deer, and thus were difficult to map.

### Diversity

We estimated genetic divergence between red deer and sika in Kintyre using the hierfstat package in R (Goudet 2005). We compared only individuals that previous analysis identified as pure species red deer or sika (McFarlane et al. 2020) and we estimated F_st_ at each individual locus following Nei (Nei 1987). We used a linear model in R (Team 2013) with Fst as the response variable, and the X chromosome as a reference to ask how the Fst of SNPs on the autosomes differed from those SNPs on the X chromosome.

### Bayesian genomic clines

We wanted to find loci with alleles that had introgressed at rates that deviated from genome wide expectations, as those alleles that move faster than expected might be under selection in the novel parental genomic background and those loci that move slower might be related to post zygotic reproductive isolation (Lexer et al. 2007). We used the program bgc (Gompert and Buerkle 2012) to estimate Bayesian genomic clines across the hybrid individuals in our population. bgc compares the genotype of each locus in each individual to that individual’s hybrid index to estimate values of α, which is comparable to a geographic cline center and β, comparable to a geographic cline slope (Gompert and Buerkle 2012).

We assigned individuals to three different populations based on their ADMIXTURE estimates and whether the credible intervals from ADMIXTURE overlapped 0 (sika) or 1 (red deer). If an individual’s credible intervals overlapped neither 0 or 1 it was considered a hybrid (McFarlane et al. 2020). Red deer and sika were each assigned to parental populations, and all admixed individuals were put into a ‘hybrid population’. This is in contrast to some previous analyses where individuals are separated based on whether they are from a population in which admixture occurs (Taylor et al. 2014, Trier et al. 2014, Royer, Streisfeld, and Smith 2016). We calculated allele frequencies for the two parental populations using PLINK (Purcell et al. 2007), while hybrid genotypes were considered individually. We ran bgc 5 independent times, for 50000 iterations each time, with a burnin of 25000 and a thinning interval of 200, and assessed convergence by eye. To be as conservative as possible when determining which loci significantly deviated from the genome wide expectation, we used the widest possible confidence intervals for each locus from the 5 chains (Janoušek et al. 2015). Loci with credible intervals that did not overlap with 0 are referred to as ‘excess’ loci. Additionally, we assumed a normal distribution for each α and β with the same mean and standard deviation as the empirical data. We then asked which SNPs had α or β estimates in the 2.5% upper and lower tails of this distribution. Those loci outside of the 95% distribution are referred to as ‘outlier loci’.

### SLiM simulations

We wanted to determine the impact of population size and history on the potential role of drift in hybridized populations. Theoretically, there is an expectation that rare, recent hybridization should result in extremely variable rates of introgression across the genome (Baird, Barton, and Etheridge 2003). We used SLiM (Haller and Messer 2017) to build some simple models that varied the rate of admixture, the length of time admixture has been occurring and the abundance ratio of each parental type population (1:1 or 3:1). We simulated 1000 individuals with a single chromosome of 1e^7^ markers, split into two populations of either 500 each or 250 and 750, and allowed both populations to evolve for 3000 generations with a standard rate of neutral mutation (0.01), typically resulting in an F_st_ between 0.40 and 0.60. Note that we did not simulate any markers to be under positive selection. We then allowed migration and interbreeding between the two populations at a given rate (0.002, 0.02, or 0.2) for a given number of generations (10, 100 or 1000). We then took the SNPs for all individuals and put them through our PLINK-ADMIXTURE-bgc pipeline (as above). One deviation from the above pipeline is that due to computational constraints bgc was only run for 2500 iterations, with a burnin of 200 iterations and a sampling interval of 2. We ran bgc 5 times for each simulation, and, as with the empirical analyses, categorized loci based on the widest possible CIs. As bgc analyses may not have converged in a such a short period of time, this could lead to wider CIs than if convergence had occurred in all chains, making this analysis conservative with respect to finding excess loci. We ran each simulation 50 times to determine what proportion of markers significantly deviated from the genome wide expectation. We did not compare to the distribution of the α and β to identify outlier loci, as this is less commonly done in the literature, and is harder to standardize across studies.

## Results

### Diversity

F_st_ varied widely among markers (Figure 1a) and across the genome (Supplementary Figure 1). While each chromosome had SNPs with F_st_ estimates that ranged from 0 to 1 (average autosomal Fst = 0.499+/0.33), the X chromosome had a higher F on average than all other chromosomes with the exception of Chromosome 25 (Figure 1b, Supplementary Table 1).

**Table 1:**
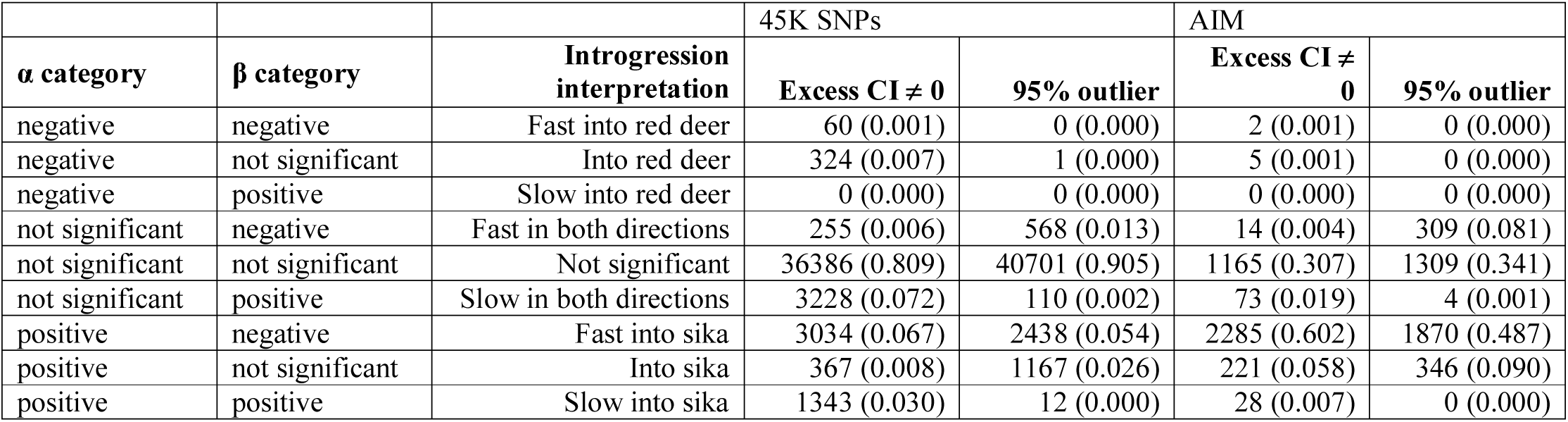
Using bgc in a red deer x sika hybrid population we categorized 44997 SNPs, and a subset of 3793 diagnostic and ancestry informative markers (AIMs) depending on the estimated center of a genomic cline (α) and rate of movement across a genomic cline (β). A SNP was considered significantly excess if the 95% confidence interval did not overlap zero, and considered an outlier if the point estimate was not within the 95% distribution for the overall genome.

**Figure 1:**
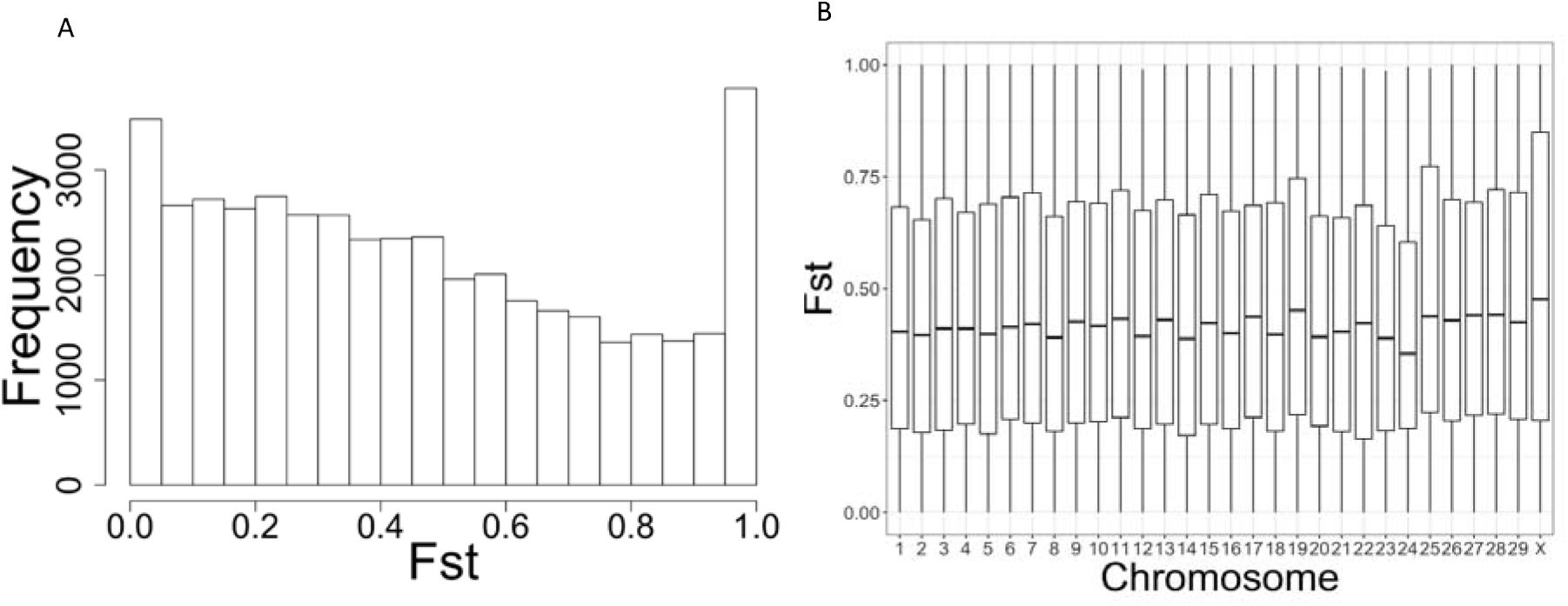
**A** Frequency of SNPs within 0.05 Fst bins, estimated using pure sika and red deer (see text). **B** Boxplot showing Fst between red deer and sika on each (bovine) chromosome. Each box shows the median, 25^th^ and 75^th^ percentile for each chromosome and each whisker extends to 5^th^ and 95^th^ percentile.

### bgc

We found substantial variation between loci in the location and rate of genomic clines between red deer and sika. Positive α can be interpreted as extreme introgression from red deer to sika, while negative α is extreme introgression from sika to red deer. While most of the 44997 SNPs that we examined were not excessively different from the genome-wide expectations there were many SNPs that were excessive compared to the genome wide expectation based on hybrid indices. Specifically, 691 (324 negative and 367 positive) SNPs were in excess for α estimates, but not for β estimates, 3483 (255 negative and 3228 positive) SNPs had β estimates that were in excess but not α estimates and 4437 other SNPs (60 negative α and β, 0 negative α and positive β, 3034 positive α and negative β, 1343 positive α and β) were in excess for both α and β (Table 1). 1168 SNPs were α outliers but not β outliers (1 negative, 1167 positive), 678 SNPs (568 negative, 110 positive) were outliers for β but not α and 2450 were outliers for both α and β (0 negative α and β, 0 negative α and positive β, 2438 positive α and negative β, 12 positive α and β). We have found substantially more excess loci with positive α estimates than negative α estimates (4744 vs 384) and substantially more positive α outliers than negative outliers (3617 vs 1). We found more positive than negative β excess SNPs (4571 vs 3349), but substantially fewer positive than negative β outlier SNPs (122 vs 3006). Excess SNPs (for either α or β) are spread across the entire genome, and occur on every chromosome (Figures 2a&b), as do outlier SNPs.

**Figure 2.**
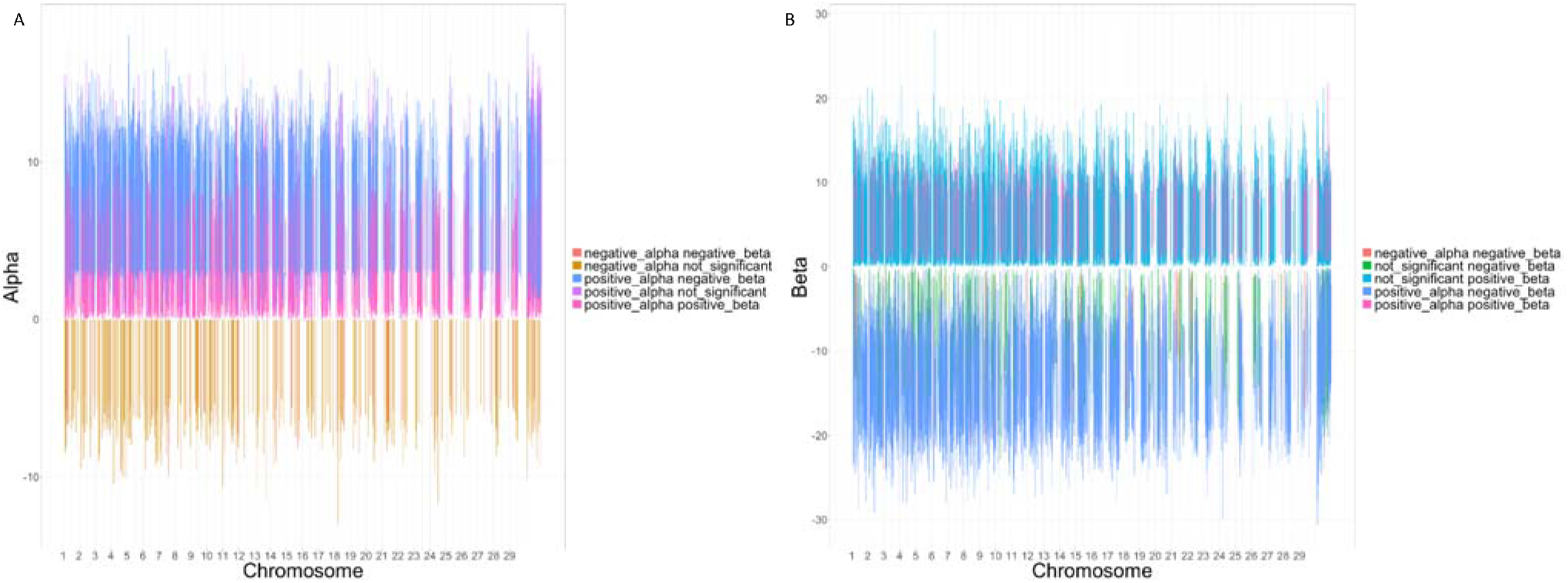
**A** – α estimates with 95% credible intervals for SNPs significantly different from zero (‘excess’), from a bgc analysis of a red deer x sika hybrid swarm in Kintyre, Scotland. α =0 can be interpreted as the genomic cline center, positive α estimates indicate alleles that are more shifted from red deer into sika than the genome wide expectation, and negative αs indicate alleles shifted from sika into red deer. **B** β estimates with 95% credible intervals for SNPs significantly different from zero (‘excess’), from a bgc analysis of a red deer x sika hybrid swarm in Kintyre, Scotland. β =0 can be interpreted as the average rate of introgression, positive β estimates are indicative of a narrow cline, and slow introgression, while negative β estimates are analogous to faster than average introgression.

When we examined only those diagnostic and ancestry informative markers we have previously identified (n=3793; McFarlane et al. 2020), we found 226 (5 negative and 221 positive) that were significantly α excess but not β excess, 87 (14 negative and 73 positive) that were significantly β excess but not α, and 2315 (2 negative α and β, 0 negative α and positive β, 2285 positive α and negative β, 28 positive α and β) that were both α and β excess. Of the AIMs, we found 346 (0 negative and 346 positive) that were α but not β outliers, 313 (309 negative and 4 positive) that were β but not α outliers and 1870 SNPs (0 negative α and β, 0 negative α and positive β, 1870 positive α and negative β, 0 positive α and β) that were significant outliers for α and β (Table 1). As was the case when we used all the SNPs, we found many more excess loci with positive α than negative α (2534 vs 7) and many more positive than negative α outlier AIM SNPs (2234 vs 0), suggesting more extreme introgression from red deer into sika than from sika into red deer. We found fewer positive than negative excess β AIM SNPs (101 vs 2301), and fewer positive than negative outlier β AIM SNPs (4 vs 2179). Similarly to when we examined all SNPs, excess and outlier α and β SNPs were found across the genome. In contrast to when we examined all SNPs, there was a substantially higher proportion of AIM SNPs that were different than the genome wide expectation (69.3% DM&AM significant excess vs 19.1% from all SNPs and 65.5% AIM significant outlier vs 9.5% from all SNPs).

### SLiM Simulations

Across the scenarios that we simulated, we found that the majority of simulated loci were not significant for either α or β estimates. However, we did find that in cases where there had only been 10 generations of admixture, and a low level of hybridization, most loci had either a positive or negative β estimate, suggesting faster or slower than expected movement through the cline (Figure 3, panels ‘sle’, ‘slo’ and ‘sme’). While the proportion of loci with significant β decreased with increasing number of generations and increased admixture, there are loci with significant β found in every other simulated scenario, with sometimes as many as 40% of loci introgressing at extreme rates when compared to the average rate of introgression across the entire genome. Additionally, in scenarios where hybridization has been progressing for longer (Figure 3, m and l rows), as many as 15% of loci have negative alpha estimates. This appears to be more extreme with increased rates of hybridization.

**Figure 3:**
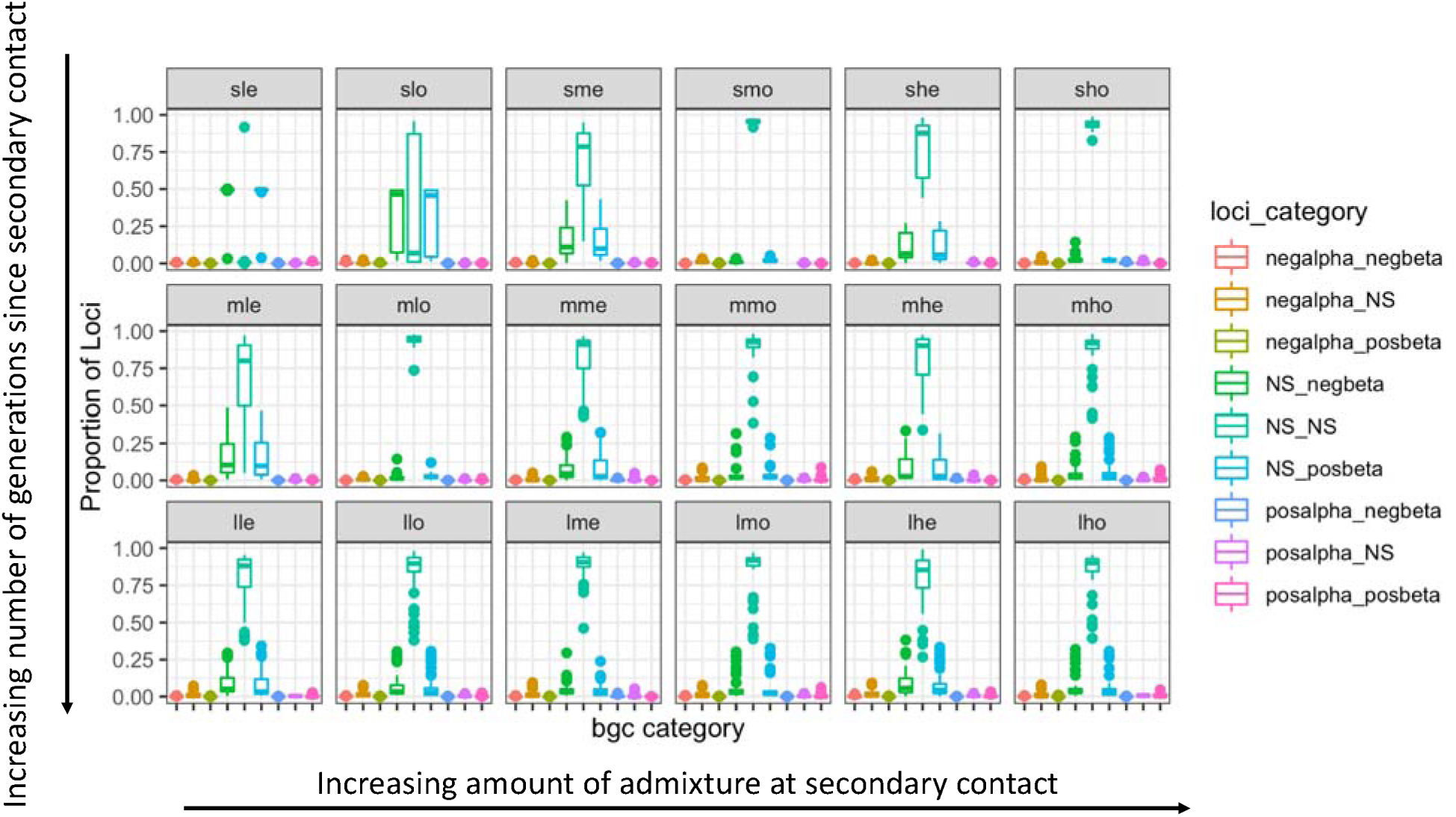
We used SLiM (Haller and Messer 2017) to simulate admixing populations that had been in secondary contact for either a short (s, 10 generations, top row), medium (m, 100 generations, middle row), or long (l, 1000 generations, bottom row) length of time since admixture started. For each length of secondary contact, we also simulated rates of migration and interbreeding between populations, as either low (l, 0.002, left two columns), medium (m, 0.02, middle two columns), or high (h, 0.2, right two columns), and the abundance ratio of each pure population, as either even (e, 1:1) or odd (o, 1:3). Each simulation was run 50 times, no selection was simulated, and we categorized (into nine categories; legend) the direction and rate of introgression among simulated hybrid individuals using bgc. Overall, introgression at most loci did not deviate from genome-wide expectation, but especially in cases with a short time since admixture started and a low rate of admixture (top, left two panels), many loci introgressed faster than genome-wide expectation despite the total absence of any selection in the simulations.

## Discussion

Using 44997 SNPs, we found extremely variable Fst between red deer and sika across all chromosomes, although the X chromosome had a substantially higher Fst than the autosomes. We also found 5128 α excess SNPs, of which 3618 are outliers and 3618 β excess SNPs of which 3128 are outliers (Table 1). When we compared these excess and outliers SNPs to our list of AIMs, we found a high proportion of AIM loci were excess and/or outliers (Table 1). This suggests that some caution should be used when interpreting the results of genomic clines of diagnostic or ancestry informative markers, as there could be a relationship between informativeness and extreme clines of these markers.

We found 4474 positive excess α SNPs (3617 outliers), and 384 negative excess α SNPs (1 outlier), which suggest cline means that have moved from red deer to sika (positive alpha) or sika to red deer more than expected based on the genomic expectation. This is in strong contrast to our simulations, which only found excess α loci in such high proportions when hybridization had been on-going for 1000 generations. Previous simulations using bgc have found substantial variation in α estimates when smaller sample sizes were simulated, even if the simulation was for only 25 generations with an admixture rate of 0.2 (Gompert and Buerkle 2011). Our empirical data set contains only 222 hybrid individuals, which is a small population compared to most of our simulations. It should be noted that the hybrid population size in our simulations varied (between approximately 45 and approximately 800), as it was a function of the admixture rate, and the stochasticity built into these individual based simulations. In any case, the 222 deer hybrids from Kintyre are substantially fewer than the 500 or 1000 hybrid individuals that were simulated in the best performing models by Gompert and Buerkle (2011). This is good reason to be cautious about interpreting excess or outlier α estimates as evidence for selection on these loci.

We found substantially more significant positive than negative excess and outlier α’s, indicating that there are more alleles that have shifted from red deer to sika than from sika to red deer. There are three possible explanations for this. First, there could be asymmetry in backcrossing, such that there is more backcrossing into sika than there is into red deer. This was previously indicated in an analysis of microsatellite data by Goodman and colleagues (Goodman et al. 1999) who estimated that the rate of backcrossing into sika was twice the rate of backcrossing into red deer (H=0.002 vs. H=0.001), although based on mitochondrial DNA, it is clear that backcrossing does proceed in both directions (Smith et al. 2018). Second, the pattern of increased positive vs. negative α estimates could be due to marker selection. The SNP chip we used was mainly designed to provide polymorphic loci for studies within red deer, and just 2250 SNPs that were selected to be diagnostic between red deer and sika (Brauning et al. 2015), although ultimately only 629 SNPs are diagnostic in our study population (McFarlane et al. 2020). These two patterns are difficult to distinguish between in our system. The sika population is less diverse than the red deer population due to a demographic history of bottlenecks and the genomic tools have been designed for use in red deer. These two processes together make it more difficult to document what could be shared alleles from sika into red deer, whereas it is easier to document the introgression of private alleles from a large, outbred, polymorphic population of red deer into sika. Further, it’s difficult to quantify the relative contribution of each of these processes to the bias that could exist. The third possible mechanism explaining the seemingly higher proportion of red deer alleles introgressing into sika than in the other direction is that, as sika are an introduced species in the UK, it is possible that some alleles that are introgressing from red deer to sika are indeed the result of adaptive introgression, because they increase the fitness of hybrids. Adaptive introgression can involve a faster response to selection in a new environment than selection on a new mutation since the allele is already proven, albeit in a different background (Hedrick 2013), and has been suggested to be a potentially positive conservation outcome of anthropogenic hybridization (Hamilton and Miller 2016). Without fitness estimates, it’s extremely difficult to demonstrate adaptive introgression in wild populations (Taylor and Larson 2019), making it difficult to tease apart these three possibilities. Empirically, we found 3349 (∼6.7%) SNPs with a negative, excess β estimate (3006 negative β outliers), suggesting that these SNPs were introgressing faster than expected between red deer and sika. While red deer and sika have been hybridizing in Scotland for at least 6-7 generations, it is possible they may have hybridized prior to introduction to Scotland, as hybridization was reported in the Irish source population before animals were introduced to Kintyre (Powerscourt 1884). Either way, this is a case of recent hybridization. The rate of backcrossing has previously been estimated using 11 microsatellite markers as 0.002 into sika and 0.001 into red deer (Goodman et al. 1999), which is consistent with our simulated ‘low’ admixture parameter. The ratio of red deer to sika is variable across Kintyre (Smith et al. 2018). Thus, our empirical work is most consistent with the ‘sle’ or the ‘slo’ simulations, where we found that most SNPs were excess β, either positive or negative (Figure 3). Thus, we found substantially fewer significant negative β SNPs than we may have expected from the simulations, highlighting that these simulations are just a toy example, rather than a highly accurate simulation of this natural system. For comparison, many studies of hybridization that have used *bgc* have not found significant β estimates. For example, a recent study of ibis hybridization using diagnostic markers found no significant negative β SNPs, in spite of the ibis hybrid zone probably only being 60 or so years old (Oswald et al. 2019). In contrast, a study of recent sole (*Solea aegyptiaca x S. senegalensis*) hybridization found 52% of all loci exhibited an extreme β value, with 26% of all loci exhibiting a negative β estimate (Souissi et al. 2018). For an example of research on an older hybrid zone, black-tailed deer and mule deer have been hybridizing for approximately 8000 years, and when genomic clines were estimated using 95 SNPs, four were found to have extreme β estimates (two positive and two negative; Haines et al. 2019). Overall, comparison of genomic cline estimates across studies and taxa is difficult, particularly given the expectation for extreme β values due to drift (Baird, Barton, and Etheridge 2003), the potential for extremely different results depending on the marker panel used (Table 1), the age of a hybrid zone, and rate of admixture between species (Figure 3). As such, a more comprehensive meta-analysis approach is likely needed to understand factors driving genomic cline variation across taxa.

Although we cannot be sure that any loci demonstrate selection in our study system we found a number of SNPs that exhibited extreme introgression as judged by α or β estimates. For example, there are 298 SNPs with Fst = 1 and a significantly negative β, suggesting that they are highly diverged between the two species, and are introgressing more quickly than would be expected in the hybrid populations. This is what we would expect if there was adaptive introgression. We didn’t find any SNPs with Fst = 1 and significantly positive β, as we might have expected to detect if there were loci with large effects on reproductive isolation. However, simulations of genomic clines that included epistatic interactions on reproductive isolation, (i.e. Bateson-Dobzhansky-Muller interactions; Dobzhansky 1937, Muller 1940) are difficult to detect using *bgc* (Gompert and Buerkle 2011), so we would not claim the lack of evidence in this case as evidence of the absence of genes involved in reproductive isolation in this system. Substantially more work is needed to address this question.

There is an expectation that when there is recent, rare hybridization, the genomic outcome of introgression is extremely stochastic (Baird, Barton, and Etheridge 2003), and it has previously been noted how difficult it is to derive a null distribution for locus-specific introgression (Gompert and Buerkle 2011). Drift can substantially increase or decrease the frequency of different blocks, in the complete absence of selection. This is consistent with what we saw in our SLiM simulations, where, when we simulated 10 generations of admixture with a rate of admixture of 0.002, we found in some cases that 50% of markers had wider clines and 50% of markers had narrower clines than predicted from the genome-wide expectation (Figure 3). As noted above, the hybrid population sizes also varied with admixture rate, particularly when hybridization was rare and had only been ongoing for 10 generations (scenarios *sle* and *slo*). This is consistent with untargeted sampling in wild populations, as, if hybridization is recent and rare, there will be proportionately fewer hybrids in the population. This confirms that extreme β estimates should not be taken as evidence of selection (Gompert and Buerkle 2012), or of adaptive introgression (Taylor and Larson 2019), as this introgression happens in the absence of selection. This is particularly true when hybridization is recent and rare, leading to relatively few hybrids in the population. Previous neutral simulations of 25 generations of admixture with an admixture rate of 0.2, comparable to our *she* and *sho* simulations but with a simulated population size of 100, found substantial variation in the estimated α or β estimates, with α being more variable than β (Gompert and Buerkle 2011). These simulations found that α or β were less variable when the population sizes simulated were 500 or 1000, although some outlier α or β loci were still found in some simulations in these cases (Gompert and Buerkle 2011). As this pattern was less extreme when hybridization had been progressing for many generations (i.e. 100 or 1000), this provides an additional rationale for researchers to quantify the length of time admixture has been occurring in their system prior to drawing conclusions (McFarlane and Pemberton 2019, Loh et al. 2013). The strength of evidence for adaptive introgression from genomic clines is, therefore, weak in more recently admixed systems, including many examples of anthropogenic hybridization. To make the case that adaptive introgression is occurring, particularly in a recent case of anthropogenic hybridization, studies must incorporate independent fitness estimates to demonstrate selection.

To conserve a species in the presence of hybridization, we must first quantify both the number of individuals in the population that are hybrids, and the proportion of alleles that could be replaced by introduced alleles, i.e. in line with the gene-based theory of conservation (Petit 2004). In our study area, we found approximately 43% of individuals are hybrids (McFarlane et al. 2020) and in the present study, we have identified 60 SNPs with both an excessive negative α and excessive negative β estimate, indicative of introgressive alleles moving from sika to red deer faster than expected. These SNPs are spread across 26 different chromosomes. Whether the pattern of these SNPs is the result of selection or drift, it is still the case that there are sika alleles that are spreading into red deer populations via hybridization faster than those at other loci. These are the genome regions that are of potential conservation concern for Scottish red deer as these alleles may most quickly replace their red deer alternates, although it should be noted that red deer are a species of least concern (IUCN 2020). Techniques such as admixture mapping could be used to try to link SNPs to phenotypes of interest (Buerkle and Lexer 2008), and then cross check these SNPs against those introgressing fastest. Such gene-targeted conservation is unlikely to be successful (Kardos and Shafer 2018), particularly since many of the traits of interest in red deer (e.g. redness, antler size and shape, size) are likely to be polygenic (Santure and Garant 2018). Specifically, body size has been found to be polygenic in a variety of taxa, including Soay sheep (Bérénos et al. 2015), bighorn sheep (Miller, Festa-Bianchet, and Coltman 2018), and polar bears (Malenfant et al. 2018). Antler shape has been found to be polygenic in Scottish red deer (Peters et al. in prep). Altogether, it seems unlikely that the 60 SNPs we have identified here would have large impacts on the phenotypic traits of interest that policy makers would seek to conserve in Scottish red deer.

Genomic clines can be used to identify loci with extreme introgression. However, genomic clines cannot be used to identify definitively alleles under selection (Gompert and Buerkle 2012, Gompert and Buerkle 2011), so different methods must be employed to distinguish between alleles undergoing adaptive introgression or involved in reproductive isolation and those loci that deviate from genomic expectations due to stochastic processes. One approach would be to study replicate hybrid zones, on the assumption that stochastic processes will act independently in each instance of secondary contact, but selection will not. Loci which have consistent excess β estimates would be the best candidates for being under selection, either for or against introgression into a novel background. In house mice, it was found that 28/41 SNPs had different genomic clines between two replicates, as assessed using a likelihood ratio test that compared the clines, encompassing both α and β, suggesting that few if any of the extreme SNPs could be related to genetic incompatibilities or adaptive introgression (Teeter et al. 2010). While it should be noted that detecting signals of even very strong selection at the genome wide level is extremely difficult, requires substantial power and a strong signal (Castro et al. 2019), those SNPs with extreme β across multiple replicate hybrid zones would be strong candidates for being involved in either adaptive introgression, or impeding gene flow between species. Future research on red deer x sika hybridization could capitalize on replicate hybrid areas across Europe (e.g. Ireland (Smith et al. 2014), Lithuania (RaŽanskė, GibieŽaitė, and Paulauskas 2017), and Poland (Biedrzycka, Solarz, and Okarma 2012)) where the many points of sika introduction have generated natural replications of this cross where selection may occur.

## Data Availability

all data and code are available at https://figshare.com/projects/Locus-specific_introgression_in_young_hybrid_swarms_drift_dominates_selection/76473

## Acknowledgements

We thank the Forestry and Land Scotland rangers, especially Fraser Robinson and Kevin McKillop for collecting samples, the Welcome Trust Clinical Research Facility Genetics Core, Edinburgh for performing the genotyping and Paul Fisher and Rudi Brauning for SNP array development. We’re also grateful to Nick Barton and Stuart Baird for discussions about the null expectation of genomic clines, as well as Alana Alexander, Zachary Gompert, Elizabeth Mandeville and Piotr Zieliński for assistance with and discussion of bgc. This project was funded by a European Research Council Advanced Grant to JMP, a Vetenskapsrådet (Swedish Research Council) International Postdoc Fellowship to SEM and Natural Environment Research Council PhD Studentships to HVS and SLS.

## Author contributions

SEM and JMP conceptualized the project, HVS and SLS collected the data, SEM analyzed the data and wrote the manuscript and all authors contributed to discussions, revised and approved the final version.

## Works Cited

Allendorf, Fred W, Robb F Leary, Paul Spruell, and John K Wenburg. 2001. “The problems with hybrids: setting conservation guidelines.” Trends in ecology & evolution 16 (11):613–622.

Allendorf, Fred W, and Gordon Luikart. 2009. Conservation and the genetics of populations: John Wiley & Sons.

Arnold, Michael L, Mark R Bulger, John M Burke, Alice L Hempel, and Joseph H Williams. 1999. “Natural hybridization: how low can you go and still be important?” Ecology 80 (2):371–381.

Baack, Eric J, and Loren H Rieseberg. 2007. “A genomic view of introgression and hybrid speciation.” Current opinion in genetics & development 17 (6):513–518.

Baird, SJE, NH Barton, and AM Etheridge. 2003. “The distribution of surviving blocks of an ancestral genome.” Theoretical population biology 64 (4):451–471.

Barton, N H, and G M Hewitt. 1985. “Analysis of hybrid zones.” Ann. Rev. Ecol. Syst. 16:113–148.

Barton, Nicholas H, and Katherine S Gale. 1993. “Genetic analysis of hybrid zones.” Hybrid zones and the evolutionary process:13–45.

Biedrzycka, Aleksandra, Wojciech Solarz, and Henryk Okarma. 2012. “Hybridization between native and introduced species of deer in Eastern Europe.” Journal of Mammalogy 93 (5):1331–1341.

Brauning, Rudiger, Paul J Fisher, Alan F McCulloch, Russell J Smithies, James F Ward, Matthew J Bixley, Cindy T Lawley, Suzanne J Rowe, and John C McEwan. 2015. “Utilization of high throughput genome sequencing technology for large scale single nucleotide polymorphism discovery in red deer and Canadian elk.” bioRxiv:027318.

Buerkle, C Alex, and Christian Lexer. 2008. “Admixture as the basis for genetic mapping.” Trends in Ecology & Evolution 23 (12):686–694.

Burri, Reto, Alexander Nater, Takeshi Kawakami, Carina F Mugal, Pall I Olason, Linnea Smeds, Alexander Suh, Ludovic Dutoit, Stanislav Bureš, and Laszlo Z Garamszegi. 2015. “Linked selection and recombination rate variation drive the evolution of the genomic landscape of differentiation across the speciation continuum of Ficedula flycatchers.” Genome Research 25 (11):1656–1665.

Bérénos, Camillo, Philip A Ellis, Jill G Pilkington, S Hong Lee, Jake Gratten, and Josephine M Pemberton. 2015. “Heterogeneity of genetic architecture of body size traits in a free-living population.” Molecular Ecology 24 (8):1810–1830.

Castro, João PL, Michelle N Yancoskie, Marta Marchini, Stefanie Belohlavy, Layla Hiramatsu, Marek Kučka, William H Beluch, Ronald Naumann, Isabella Skuplik, and John Cobb. 2019. “An integrative genomic analysis of the Longshanks selection experiment for longer limbs in mice.” elife 8:e42014.

Charlesworth, Brian. 1998. “Measures of divergence between populations and the effect of forces that reduce variability.” Molecular Biology and Evolution 15 (5):538–543.

Cruickshank, Tami E, and Matthew W Hahn. 2014. “Reanalysis suggests that genomic islands of speciation are due to reduced diversity, not reduced gene flow.” Molecular Ecology 23 (13):3133–3157.

Dobzhansky, T. 1937. “Genetics and the origin of species.” 374.

Gompert, Z, and CA Buerkle. 2012. “bgc: software for Bayesian estimation of genomic clines.” Molecular Ecology Resources 12 (6):1168–1176.

Gompert, Zachariah, and C Alex Buerkle. 2009. “A powerful regression-based method for admixture mapping of isolation across the genome of hybrids.” Molecular Ecology 18 (6):1207–1224.

Gompert, Zachariah, and C Alex Buerkle. 2011. “Bayesian estimation of genomic clines.” Molecular Ecology 20 (10):2111–2127.

Goodman, Simon J, Nick H Barton, Graeme Swanson, Kate Abernethy, and Josephine M Pemberton. 1999. “Introgression through rare hybridization: a genetic study of a hybrid zone between red and sika deer (genus Cervus) in Argyll, Scotland.” Genetics 152 (1):355–371.

Goudet, Jérôme. 2005. “Hierfstat, a package for R to compute and test hierarchical F-statistics.” Molecular Ecology Notes 5 (1):184–186.

Grabenstein, Kathryn C, and Scott A Taylor. 2018. “Breaking Barriers: Causes, Consequences, and Experimental Utility of Human-Mediated Hybridization.” Trends in Ecology & Evolution.

Haines, Margaret L, Gordon Luikart, Stephen J Amish, Seth Smith, and Emily K Latch. 2019. “Evidence for adaptive introgression of exons across a hybrid swarm in deer.” BMC Evolutionary BNiology 19 (1):199.

Haller, Benjamin C, and Philipp W Messer. 2017. “SLiM 2: Flexible, interactive forward genetic simulations.” Molecular Biology and Evolution 34 (1):230–240.

Hamilton, Jill A, and Joshua M Miller. 2016. “Adaptive introgression as a resource for management and genetic conservation in a changing climate.” Conservation Biology 30 (1):33–41.

Hedrick, Philip W. 2013. “Adaptive introgression in animals: examples and comparison to new mutation and standing variation as sources of adaptive variation.” Molecular Ecology 22 (18):4606–4618.

Hewitt, Godfrey M. 1988. “Hybrid zones-natural laboratories for evolutionary studies.” Trends in Ecology & Evolution 3 (7):158–167.

Huisman, Jisca, Loeske EB Kruuk, Philip A Ellis, Tim Clutton-Brock, and Josephine M Pemberton. 2016. “Inbreeding depression across the lifespan in a wild mammal population.” Proceedings of the National Academy of Sciences 113 (13):3585–3590.

IUCN. 2020. IUCN Red List of Threatened Species. Version 2020.1. <www.iucnredlist.org>.

Janoušek, Václav, Pavel Munclinger, Liuyang Wang, Katherine C Teeter, and Priscilla K Tucker. 2015. “Functional organization of the genome may shape the species boundary in the house mouse.” Molecular Biology and Evolution 32 (5):1208–1220.

Johnston, Susan E, Jisca Huisman, Philip A Ellis, and Josephine M Pemberton. 2017. “A high-density linkage map reveals sexually-dimorphic recombination landscapes in red deer (Cervus elaphus).” G3: Genes, Genomes, Genetics 8 (7):2265–2276.

Kardos, Marty, and Aaron BA Shafer. 2018. “The peril of gene-targeted conservation.” Trends in Ecology & Evolution 33 (11):827–839.

Lexer, C, CA Buerkle, JA Joseph, B Heinze, and MF Fay. 2007. “Admixture in European Populus hybrid zones makes feasible the mapping of loci that contribute to reproductive isolation and trait differences.” Heredity 98 (2):74–84.

Loh, Po-Ru, Mark Lipson, Nick Patterson, Priya Moorjani, Joseph K Pickrell, David Reich, and Bonnie Berger. 2013. “Inferring admixture histories of human populations using linkage disequilibrium.” Genetics 193 (4):1233–1254.

Malenfant, René M, Corey S Davis, Evan S Richardson, Nicholas J Lunn, and David W Coltman. 2018. “Heritability of body size in the polar bears of Western Hudson Bay.” Molecular Ecology Resources 18 (4):854–866.

Mallet, James, Nick Barton, Gerard Lamas, Jose Santisteban, Manuel Muedas, and H Eeley. 1990. “Estimates of selection and gene flow from measures of cline width and linkage disequilibrium in Heliconius hybrid zones.” Genetics 124 (4):921–936.

McFarlane, S Eryn, Darren C Hunter, Helen V Senn, Stephanie L Smith, Rebecca Holland, Jisca Huisman, and Josephine M Pemberton. 2020. “Increased genetic marker density reveals high levels of admixture between red deer and introduced Japanese sika in Kintyre, Scotland.” Evolutionary Applications 13 (2):432–441.

McFarlane, S Eryn, and Josephine M Pemberton. 2019. “Detecting the true extent of introgression during anthropogenic hybridization.” Trends in Ecology & Evolution 34 (4):315–326.

McFarlane, S. Eryn, elen V Senn, Stephanie L Smith and Josephine M Pemberton. 2020. Locus-specific introgression in young hybrid swarms: drift dominates selection – dataset. https://figshare.com/projects/Locus-specific_introgression_in_young_hybrid_swarms_drift_dominates_selection/76473

Miller, Joshua M, Marco Festa-Bianchet, and David W Coltman. 2018. “Genomic analysis of morphometric traits in bighorn sheep using the Ovine Infinium® HD SNP BeadChip.” PeerJ 6:e4364.

Muller, Hermann J. 1940. “Bearing of the Drosophila work on systematics.” The New Systematics:185–268.

Nei, Masatoshi. 1987. Molecular Evolutionary Genetics: Columbia university press. Oswald,

Jessica A, Michael G Harvey, Rosalind C Remsen, DePaul U Foxworth, Donna L Dittmann, Steven W Cardiff, and Robb T Brumfield. 2019. “Evolutionary dynamics of hybridization and introgression following the recent colonization of Glossy Ibis (Aves: Plegadis falcinellus) into the New World.” Molecular Ecology 28 (7):1675–1691.

Parchman, TL, Z Gompert, Michael J Braun, RT Brumfield, DB McDonald, JAC Uy, G Zhang, ED Jarvis, BA Schlinger, and CA Buerkle. 2013. “The genomic consequences of adaptive divergence and reproductive isolation between species of manakins.” Molecular Ecology 22 (12):3304–3317.

Parmesan, Camille, and Gary Yohe. 2003. “A globally coherent fingerprint of climate change impacts across natural systems.” Nature 421 (6918):37–42.

Peters, Lucy, Jisca Huisman, Loeske EB Kruuk, Josephine M Pemberton, and Susan E Johnston. in prep. “Antler morphology has a polygenic genetic architecture in wild red deer (Cervus elaphus).”

Petit, Rémy J. 2004. “Biological invasions at the gene level.” Diversity and Distributions 10 (3):159–165.

Powerscourt, Viscount. 1884. “On the Acclimatization of the Japanese Deer at Powerscourt.” Proceedings of the Zoological Society of London:207–209.

Pulido-Santacruz, Paola, Alexandre Aleixo, and Jason T Weir. 2018. “Morphologically cryptic Amazonian bird species pairs exhibit strong postzygotic reproductive isolation.” Proceedings of the Royal Society B: Biological Sciences 285 (1874):20172081.

Purcell, S, B Neale, K Todd-Brown, L Thomas, MAR Ferreira, D Bender, J Maller, P Sklar, PIW de Bakker, MJ Daly, and PC Sham. 2007. “PLINK: a toolset for whole-genome association and population-based linkage analysis.” American Journal of Human Genetics 81.

Ratcliffe, PR. 1987. “Distribution and current status of sika deer, Cervus nippon, in Great Britain.” Mammal Review 17 (1):39–58.

RaŽanskė, Irma, Justina Monika Gibiežaitė, and Algimantas Paulauskas. 2017. “Genetic analysis of red deer (Cervus elaphus) and sika deer (Cervus nippon) to evaluate possible hybridisation in Lithuania.” Baltic forestry. Girionys: Lietuvos mišku institutas, 2017, vol. 23, no. 3.

Rhymer, Judith M, and Daniel Simberloff. 1996. “Extinction by hybridization and introgression.” Annual Review of Ecology and Systematics:83–109.

Royer, Anne M, Matthew A Streisfeld, and Christopher Irwin Smith. 2016. “Population genomics of divergence within an obligate pollination mutualism: Selection maintains differences between Joshua tree species.” American Journal of Botany 103 (10):1730–1741.

Santure, Anna W, and Dany Garant. 2018. “Wild GWAS—association mapping in natural populations.” Molecular Ecology Resources 18 (4):729–738.

Scottish Wildlife Trust. 2013. https://scottishwildlifetrust.org.uk/news/can-you-spot-all-of-scotlands-big-5/#:~:text='Scotland's%20Big%205'%20celebrates%20the,animals%20in%20their%20natural%20habitat. Accessed 16.09.2020.

Senn, Helen V, Nick H Barton, Simon J Goodman, GM Swanson, KA Abernethy, and Josephine M Pemberton. 2010. “Investigating temporal changes in hybridization and introgression in a predominantly bimodal hybridizing population of invasive sika (Cervus nippon) and native red deer (C. elaphus) on the Kintyre Peninsula, Scotland.” Molecular Ecology 19 (5):910–924.

Senn, Helen V, and Josephine M Pemberton. 2009. “Variable extent of hybridization between invasive sika (Cervus nippon) and native red deer (C. elaphus) in a small geographical area.” Molecular Ecology 18 (5):862–876.

Senn, Helen V, Graeme M Swanson, Simon J Goodman, Nicholas H Barton, and Josephine M Pemberton. 2010. “Phenotypic correlates of hybridisation between red and sika deer (genus Cervus).” Journal of Animal Ecology 79 (2):414–425.

Smith, Stephanie L, Ruth F Carden, Barry Coad, Timothy Birkitt, and Josephine M Pemberton. 2014. “A survey of the hybridisation status of Cervus deer species on the island of Ireland.” Conservation Genetics 15 (4):823–835.

Smith, Stephanie L, Helen V Senn, Sílvia Pérez-Espona, Megan T Wyman, Elizabeth Heap, and Josephine M Pemberton. 2018. “Introgression of exotic Cervus (nippon and canadensis) into red deer (Cervus elaphus) populations in Scotland and the English Lake District.” Ecology and Evolution 8 (4):2122–2134.

Souissi, Ahmed, François Bonhomme, Manuel Manchado, Lilia Bahri-Sfar, and Pierre-Alexandre Gagnaire. 2018. “Genomic and geographic footprints of differential introgression between two divergent fish species (Solea spp.).” Heredity 121 (6):579–593.

Sung, Cheng-Jung, Katherine L Bell, Chris C Nice, and Noland H Martin. 2018. “Integrating Bayesian genomic cline analyses and association mapping of morphological and ecological traits to dissect reproductive isolation and introgression in a Louisiana Iris hybrid zone.” Molecular Ecology 27 (4):959–978.

Taylor, Scott A, Robert L Curry, Thomas A White, Valentina Ferretti, and Irby Lovette. 2014. “Spatiotemporally consistent genomic signatures of reproductive isolation in a moving hybrid zone.” Evolution 68 (11):3066–3081.

Taylor, Scott A, and Erica L Larson. 2019. “Insights from genomes into the evolutionary importance and prevalence of hybridization in nature.” Nature Ecology & Evolution 3 (2):170–177.

Team, R Core. 2013. “R: A language and environment for statistical computing.” R Foundation for Statistical Computing, Vienna, Austria. URL http://www.R-project.org/.

Teeter, Katherine C, Lisa M Thibodeau, Zachariah Gompert, C Alex Buerkle, Michael W Nachman, and Priscilla K Tucker. 2010. “The variable genomic architecture of isolation between hybridizing species of house mice.” Evolution: International Journal of Organic Evolution 64 (2):472–485.

Todesco, Marco, Mariana A Pascual, Gregory L Owens, Katherine L Ostevik, Brook T Moyers, Sariel Hübner, Sylvia M Heredia, Min A Hahn, Celine Caseys, and Dan G Bock. 2016. “Hybridization and extinction.” Evolutionary Applications 9 (7):892–908.

Trier, Cassandra N, Jo S Hermansen, Glenn-Peter Sætre, and Richard I Bailey. 2014. “Evidence for mito-nuclear and sex-linked reproductive barriers between the hybrid Italian sparrow and its parent species.” PLoS Genetics 10 (1):e1004075.

