## supplementary files for "Locus-specific introgression in young hybrid swarms: drift dominates selection"

**Supplementary Material**

Supplementary Table S1: Comparison of Fst on the X chromosome to other chromosomes. SNPs on the X chromosome have significantly higher Fst's than SNPs on all the autosomes with the exception of chromosome 25.

| Chromosome | Estimate | Std. Error | t value | p value |
|---|---|---|---|---|
| (Intercept) | 0.510 | 0.01 | 81.784 | < 2.00E-16 |
| 1 | -0.064 | 0.01 | -7.607 | 2.86E-14 |
| 2 | -0.079 | 0.01 | -9.024 | < 2.00E-16 |
| 3 | -0.059 | 0.01 | -6.457 | 1.08E-10 |
| 4 | -0.067 | 0.01 | -7.498 | 6.58E-14 |
| 5 | -0.068 | 0.01 | -7.552 | 4.37E-14 |
| 6 | -0.053 | 0.01 | -5.85 | 4.96E-09 |
| 7 | -0.050 | 0.01 | -5.419 | 6.02E-08 |
| 8 | -0.075 | 0.01 | -8.145 | 3.89E-16 |
| 9 | -0.052 | 0.01 | -5.501 | 3.79E-08 |
| 10 | -0.061 | 0.01 | -6.358 | 2.06E-10 |
| 11 | -0.039 | 0.01 | -4.161 | 3.18E-05 |
| 12 | -0.074 | 0.01 | -7.31 | 2.71E-13 |
| 13 | -0.052 | 0.01 | -5.149 | 2.63E-07 |
| 14 | -0.079 | 0.01 | -7.817 | 5.54E-15 |
| 15 | -0.051 | 0.01 | -5.025 | 5.06E-07 |
| 16 | -0.073 | 0.01 | -7.084 | 1.42E-12 |
| 17 | -0.048 | 0.01 | -4.656 | 3.23E-06 |
| 18 | -0.063 | 0.01 | -5.724 | 1.05E-08 |
| 19 | -0.027 | 0.01 | -2.427 | 0.015245 |
| 20 | -0.070 | 0.01 | -6.641 | 3.15E-11 |
| 21 | -0.073 | 0.01 | -6.728 | 1.74E-11 |
| 22 | -0.066 | 0.01 | -5.78 | 7.52E-09 |
| 23 | -0.079 | 0.01 | -6.338 | 2.36E-10 |
| 24 | -0.104 | 0.01 | -9.235 | < 2.00E-16 |
| 25 | -0.021 | 0.01 | -1.612 | 0.106877 |
| 26 | -0.050 | 0.01 | -4.135 | 3.56E-05 |
| 27 | -0.047 | 0.01 | -3.581 | 0.000343 |
| 28 | -0.037 | 0.01 | -2.915 | 0.00356 |
| 29 | -0.045 | 0.01 | -3.626 | 0.000289 |

Supplementary Figure 1: We calculated the $F_{st}$ between red deer and sika on the Kintyre peninsula using 44997 SNPs. We have plotted $F_{st}$ across the map position of each chromosome, including the X chromosome. We used the bovine map positions and linkage map because many diagnostic and ancestry informative markers, which were not polymorphic in sika, were not mapped on the *Cervus* linkage map (Johnston et al. 2017). For this reason, we present only 29 autosomes, as cattle have 29 autosomes, although red deer have 33. Map positions have been constrained between 0 and 1 for graphical purposes only.

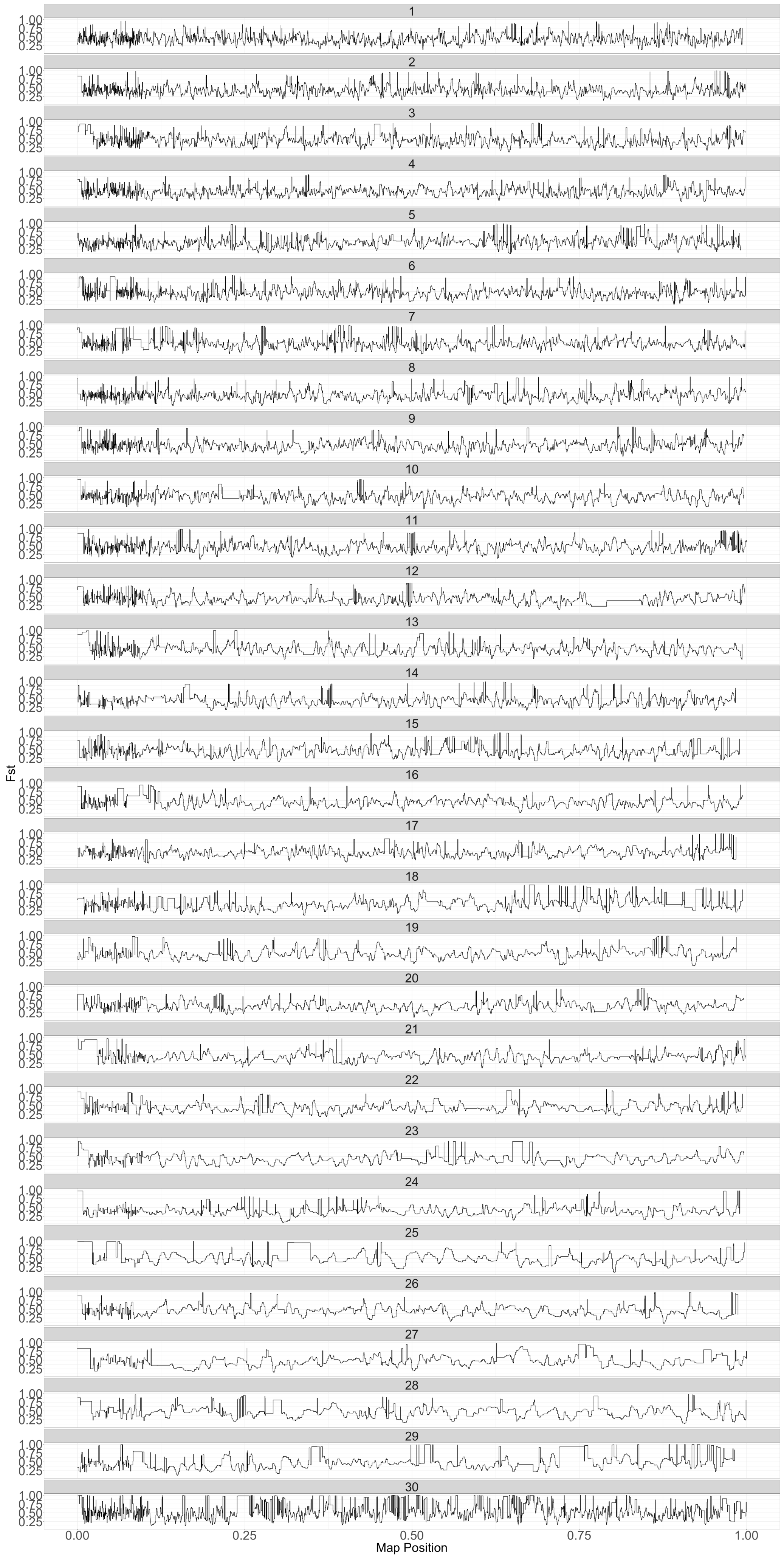

Fst
1.00
0.75
0.50
0.25
0.00
0.25
0.50
0.75
1.00
Map Position
